# Decoding menopause-induced tissue fibrosis using pan-tissue network inference

**DOI:** 10.1101/2025.07.03.663094

**Authors:** Hirotaka Iijima, Atsushi Yamashita, Jenna L Galloway, Nam Vo, Hak Soo Choi, Fabrisia Ambrosio

## Abstract

Menopause drives fibrotic remodeling and consequent tissue dysfunction across multiple organs, yet the tissue-conserved versus tissue-specific mechanisms underlying this phenomenon remain poorly defined. Here, we employed a network inference framework to uncover how menopause triggers coordinated shifts in intercellular signaling cascades that promote fibrosis. We leveraged publicly archived single-cell RNA-sequencing data from the liver, lung, pancreas, and skeletal muscle of ovariectomized and control mice. Given the central role of immune cells in orchestrating inflammation and fibrosis, we focused our analysis on immune cell populations. Using canonical markers of immune cells, we annotated major cell types and reconstructed cell–cell interaction networks to map transcriptional responses to hormonal shifts. Network analysis revealed pervasive reshaping of intercellular signaling common to all tissues evaluated in the setting of menopause. Within this immune-centered framework, we found that estrogen-responsive macrophages consistently function as major signaling hubs across all tissues evaluated, exhibiting extensive interactions with myofibroblasts—key drivers of extracellular matrix production and fibrotic remodeling. Notably, these shared signaling patterns were not detectable using conventional differential gene expression analysis, which revealed minimal overlap in gene-level responses in macrophages across tissues. In addition to conserved patterns, we identified tissue-specific interaction networks that reflect unique immune adaptations to hormonal loss. As an example, natural killer cells acted as a signaling hub in muscle-specific patterns, suggesting their direct contribution to menopausal skeletal muscle adaptation. Tissue-specific patterning was also evident in the liver, lung, and pancreas, where other immune cell types, such as CD8^+^ T cells and endothelial cells, functioned as prominent signaling hubs, indicating diverse remodeling of the immune microenvironment. The network approach introduced here represents a systems-level framework for mapping multicellular network rewiring following hormonal depletion and highlights conserved immune–stromal modules as potential therapeutic targets to prevent menopause-associated dysfunction.

## INTRODUCTION

Hormonal shifts accompanying menopause profoundly disrupt tissue homeostasis across multiple organ systems, leading to functional decline and increased susceptibility to pathology in older women.^1,2^ These changes appear to involve systemic immune remodeling, including shifts in both innate and adaptive immune cell populations^,3,4^ contributing to a state of chronic, low-grade inflammation—commonly referred to as “*inflammaging*”. This sustained inflammatory state, in turn, fosters a pro-fibrotic tissue environment.^5^ A fibrotic environment is characterized by sustained fibroblast activation, aberrant extracellular matrix remodeling, and impaired tissue regeneration, all of which can be regulated by immune cell signals.^6^ Immune-driven fibrosis is a hallmark of chronic diseases that disproportionately affect postmenopausal women—including cardiovascular disease, sarcopenia, and pulmonary fibrosis.^7^ Despite these strong associations, the mechanisms linking menopause-induced immune shifts to aberrant remodeling remain largely undefined.

To effectively disentangle the integrated and dynamic crosstalk between hormonal cues, immune cell behavior, and tissue-specific microenvironments, a comprehensive understanding of how distinct immune cell types respond to hormonal shifts in both conserved and tissue-specific manner is needed. Such an understanding requires a mechanistic framework capable of capturing cell–cell interactions and context-dependent gene regulation. Single-cell transcriptomics, combined with network inference, offers a powerful means to map how menopause reconfigures the immune regulatory landscape both within individual cell types and across cellular ecosystems. Network inference, in particular, enables the reconstruction of regulatory relationships between genes and cells based on expression data, effectively revealing “*who is talking to whom*” within the system.^8^ By translating these interactions into networks, we can discover master regulators and signaling nodes that orchestrate pathological processes such as fibrosis.^9^ This systems-level approach provides critical insights into the coordinated transcriptional programs and intercellular signaling circuits that drive tissue dysfunction in the postmenopausal state.

Here, we employed a network inference framework to capture tissue-specific transcriptional programs and regulatory hubs at single-cell resolution. By applying cell–cell interaction network analysis to archived single-cell RNA-sequencing data from ovariectomized mice, we identified both conserved and tissue-specific immune–stromal circuits involved in fibrotic tissue remodeling. Leveraging archived data obtained from various organs from a murine model of surgical menopause, we constructed an integrated network of coordinated transcriptional programs that represent tissue-level responses to sex hormone deprivation. The network-driven approach employed here offers a powerful platform for the mechanistic dissection of menopause-induced cellular dysfunction and provides a foundation to identify both convergent and distinct therapeutic targets that transcend individual organ systems.

## RESULTS

### Menopause reshapes the immune landscape

Fibrosis is a common pathological feature of menopause across multiple tissues.^10^ Given emerging evidence has implicated immune dysregulation as a key driver of fibrotic progression,^11^ as the first step of our investigation, we hypothesized that menopause induces alterations in immune cell composition that may underlie the widespread fibrotic remodeling. Specifically, this initial analysis focused on identifying shifts in immune cell populations that are common to all tissues evaluated.

To test our initial hypothesis, we analyzed archived single-cell RNA sequencing (scRNA-seq) data from C57BL/6 young female mice subjected to ovariectomy (OVX) as a widely used model of surgically-induced menopause.^12^ We focused on four organs—liver, lung, pancreas, and skeletal muscle—each of which has been previously identified as a site of menopause-associated aberrant tissue remodeling.^11,13-15^ All four tissue samples were collected from the same groups of OVX or sham-operated mice in a single study, thus reducing variability and enabling direct tissue comparisons. Data preprocessing was conducted using the established Seurat pipeline, including quality control, normalization, and dimensionality reduction (**Figure 1A**).^16^ Immune cell annotation was performed using the ScType pipeline, ensuring robust classification of immune subpopulations across all tissue datasets.^17^

**Figure 1.**
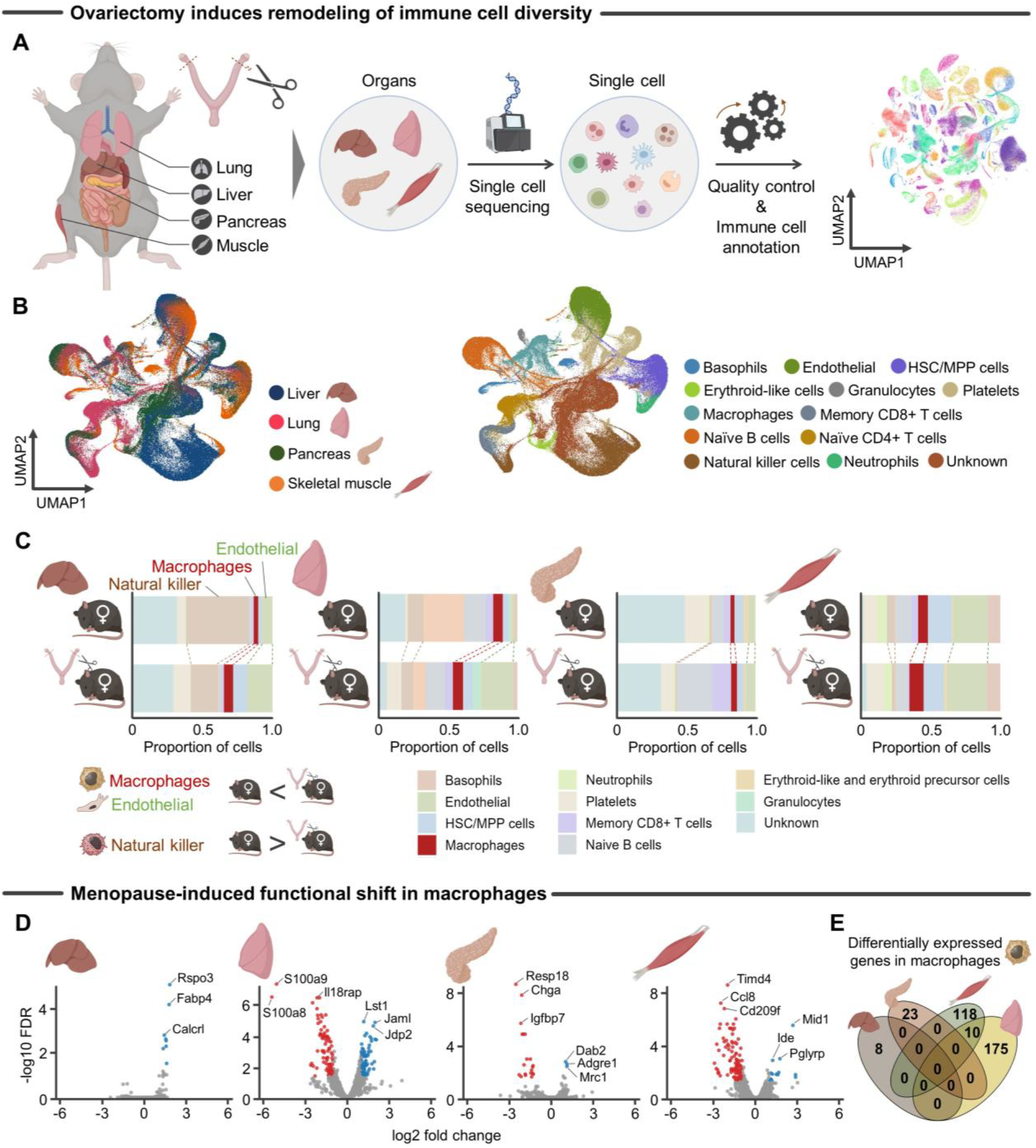
Menopause drives immune cell expansion and functional shifts. **A**, Schematic representation of analytical flow for single cell RNA-seq, including immune cell annotation from control and ovariectomized female mice across four tissues (liver, lung, pancreas, and skeletal muscle). **B**, UMAP visualization of the immune cell compartment, with cells colored by their tissue of origin or predicted immune cell types as determined by scType.^17^ **C**, UMAP visualization of the immune cell compartment, with cells colored according to menopausal status (control vs. OVX). **D**, Volcano plot highlighting the functional shift of macrophages in response to menopausal hormonal shift (control vs. OVX). Differential expression analysis was performed in “pseudo-bulk” samples macrophages. E, Venn diagram showing that differentially expressed genes in macrophages exhibited minimal overlap across tissues when comparing control vs. OVX. *Abbreviation: HSC/MPP, hematopoietic stem and multipotent progenitor; UMAP, Uniform manifold approximation and projection; OVX, ovariectomized*.

Uniform Manifold Approximation and Projection (UMAP) analysis of scRNA-seq data revealed a core set of 12 major immune cell types consistently present across all tissues (**Figure 1B**). Building on these observations, we uncovered a menopause-driven shift in immune cell diversity across multiple organs (**Figure 1C**). Notably, OVX mice exhibited an expansion of several immune cell subsets, with increased proportions of macrophages and endothelial cells observed in multiple tissues, although the magnitude of change varied by organ. For example, macrophage levels remained relatively unchanged in the lung, while increases were more evident in the liver and skeletal muscle. In contrast, natural killer cells tended to decline in several tissues, although this trend was not prominent in the pancreas, where natural killer cell proportions remained relatively stable (**Figure 1C**).

To further explore the consequences of these shifts, we examined the transcriptional landscape of macrophages, one of the most prominently altered populations. Analysis of a “pseudo-bulk” dataset of aggregated macrophages from each sample revealed menopause-associated transcriptional changes (**Figure 1D**). Interestingly, differential gene expression analysis of the macrophage population showed that these changes were highly tissue-specific, with minimal overlap in differentially expressed genes across the four organs examined (**Figure 1E**). While previous studies have primarily focused on immune changes within the female reproductive tract or individual organs,^3,4^,18 our analysis demonstrates that menopause-driven remodeling of tissue-resident immune cells is a systemic phenomenon spanning multiple organ systems.

### Deconstructing paracrine signaling cascades in menopausal tissue remodeling

Traditional single-cell clustering approaches often fail to capture the dynamic intercellular signaling underlying tissue responses to menopause. Methods such as UMAP-based clustering are effective for identifying discrete cellular states but fall short in capturing the coordinated nature of cell-cell crosstalk. This limitation is particularly consequential in the context of menopause, where hormonal cues initiate signaling cascades that extend beyond individual cell types. While hormone receptor signaling in target cells has been extensively characterized, downstream paracrine communication—critical for orchestrating multicellular responses such as tissue maintenance and remodeling—remains poorly resolved at the resolution of single cells (**Figure 2A**). Systemic hormonal shifts, as occurs with OVX, likely disrupt these paracrine networks, altering intercellular interactions in a way that is not readily captured by conventional clustering methods. Similar limitations have been observed in studies of the human breast, where hormone-driven tissue remodeling is mediated predominantly through indirect paracrine effects that are often overlooked by state-based single-cell analyses.^19^ Consistently, our differential expression analysis of macrophages showed minimal overlap in gene changes across tissues (**Figure 1E**).

**Figure 2.**
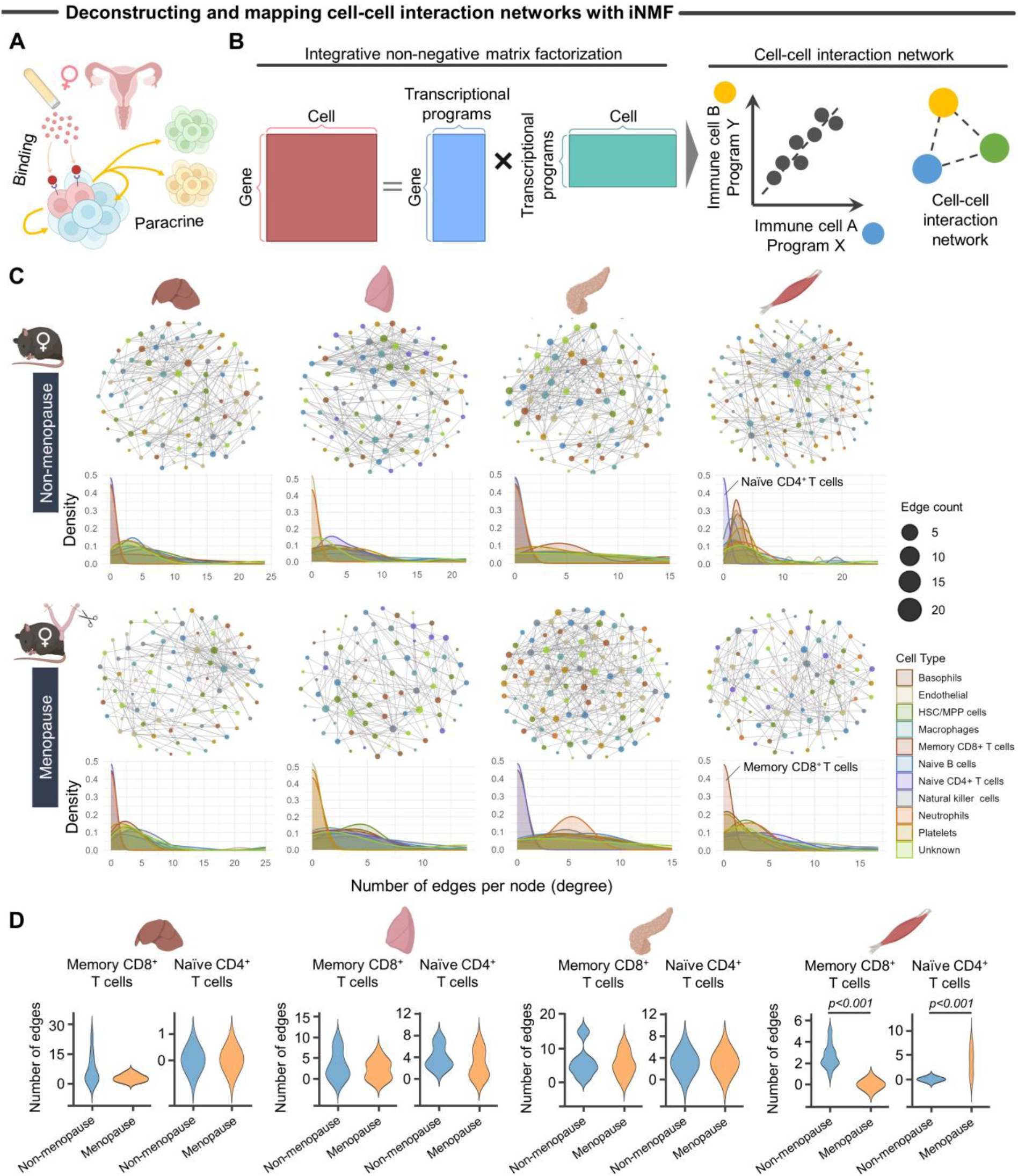
Menopause orchestrates dynamic cellular crosstalk in tissue remodeling. **A**, Schematic representation of paracrine signaling disrupted by hormone depletion. Circulating hormones bind to receptors on target cells, activating transcriptional programs that release paracrine factors to coordinate neighboring cells. OVX-induced hormone loss impairs this cascade, potentially disrupting tissue homeostasis and remodeling. **B**, Schematic representation of the cell-cell interaction network. To identify transcriptional programs, we applied iNMF to each immune cell type. Pairwise correlations between cell transcriptional programs were used to construct a tissue-level map of cell-cell interactions. **C**, Network graph illustrating correlated transcriptional programs (i.e., cell-cell interactions) in each tissue based on menopausal status. Nodes represent distinct transcriptional programs in specific immune cell types, while edges indicate significant correlations between programs (Pearson correlation coefficient >0, *p* <0.25). **D**, Menopause significantly changed connectivity and transcriptional activity of memory CD8^+^ and naïve CD4^+^ T cell populations in a tissue-dependent manner. Statistical analyses were performed using a two-tailed Student *t*-test. *Abbreviation: HSC/MPP, hematopoietic stem and multipotent progenitor; iNMF, integrative non-negative matrix factorization; OVX, ovariectomized*.

To address this gap, we used integrative non-negative matrix factorization (iNMF) to reconstruct cell-cell interaction networks, as per established methods.^19^ Non-negative matrix factorization (NMF) is a dimensionality reduction technique that factorizes a large gene expression matrix into two smaller matrices with only non-negative values.^20^ This non-negativity constraint is particularly suitable for transcriptomic data, where gene expression levels are inherently non-negative and additive. As a result, NMF enables biologically intuitive interpretation, identifying co-expressed gene programs that contribute in a compositional manner. iNMF extends traditional NMF by jointly analyzing multiple single-cell datasets, allowing us to extract lower-dimensional features that capture both shared and unique transcriptional signatures^.21^

**Figure 2B** illustrates the underlying theory, wherein iNMF factorizes the aggregated single-cell gene expression matrix into two non-negative matrices: one representing transcriptional programs (i.e., groups of genes that are co-expressed), while the other describes metacells (i.e., reflecting how strongly each cell expresses each transcriptional program). These transcriptional programs are biologically interpretable and correspond to latent signaling pathways that are often shared across multiple cell types but may differ in their relative activity or expression patterns. By leveraging differences in transcriptional profiles and cell-type composition across samples, iNMF enables us to identify coordinated patterns of intercellular communication that would be challenging, if not impossible, to detect using conventional approaches. Specifically, this framework allows the reconstruction of cell-cell interaction networks based on shared transcriptional programs, providing a systems-level perspective on multicellular organization and communication.

Building on this foundation, we applied iNMF to scRNA-seq datasets from four organs across non-menopausal and menopausal groups to construct the cell–cell interaction network (**Figure 2C**). The resulting network displayed scale-free architecture, a hallmark of biological systems, characterized by a few highly connected hubs that represent key cells or transcriptional programs, while the majority of nodes display only a limited number of connections. This network topology implies that tissue homeostasis is shaped by a subset of influential cell interactions rather than uniform contributions from all cell types. Notably, we found that the scale-free properties of the network varied according to tissue type and menopause status (**Figure 2C**). To clearly illustrate this, we highlight skeletal muscle as a representative example, where we observed a rewiring of interactions involving CD8^+^ and CD4^+^ T cells (**Figure 2D**). In contrast, such alterations in T cell balance were not evident in other tissues examined, including liver, lung, and pancreas (**Figure 2D**). The altered CD4+/CD8+ T cell balance—marked by increased CD8+ T cell exhaustion—may reflect immune remodeling that impairs regenerative capacity in menopausal muscle^.22^

### Cross-tissue coordination of cell-cell interactions in menopausal remodeling

Building on the scale-free structure identified in our cell-cell interaction network **(Figure 2C**), we next investigated whether these signaling patterns reflect a coordinated cross-tissue response or organ-specific adaptations. To do this, we developed a “*differential*” cell-cell interaction analysis framework within our iNMF-based approach to systematically compare signaling dynamics across the four menopausal tissues of interest (**Figure 3A**). This method allowed us to distinguish tissue-conserved from tissue-specific adaptations by identifying pairs of cell types with specific transcriptional programs that exhibit consistent shifts following menopause.

**Figure 3.**
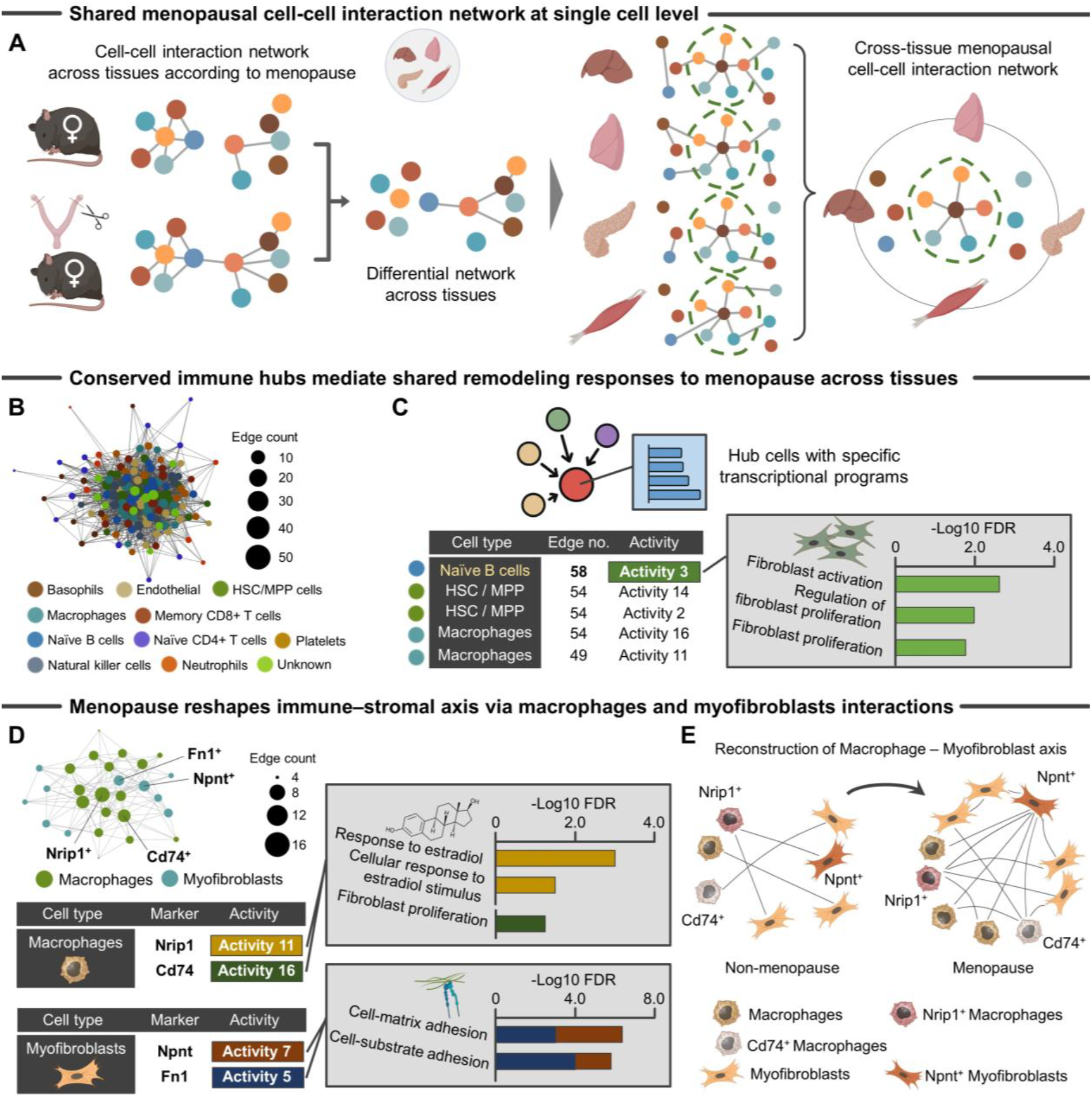
Cross-tissue immune network remodeling in menopause. **A**, Schematic representation of the differential cell-cell interaction analysis framework developed to identify menopause-associated *conserved* shifts in transcriptionally defined cell-cell signaling across four tissues. **B**, Network visualization of shared menopausal cell-cell interaction network reveals conserved remodeling patterns, including the redistribution of immune cell subsets and alterations in signaling hierarchies. Nodes represent distinct transcriptional programs in specific immune cell types, while edges indicate significant correlations between programs across tissues (Pearson correlation coefficient >0, *p* <0.25). Menopause-associated changes in immune system diversity highlight a conserved response rather than isolated, tissue-specific effects. **C**, Conserved immune hubs, particularly subsets of B cells and macrophages, emerge as central coordinators of menopausal shift. Naïve B cells exhibit transcriptional programs associated with fibroblast activation and proliferation. **D**, Network representation of menopause-specific interactions between macrophages and myofibroblast. Estrogen-responsive Nrip1^+^ macrophages and fibroblast proliferation–associated Cd74^+^ macrophages act as central hubs in the immune-stromal signaling axis, establishing robust interactions with Npnt^+^ or Fn1^+^ myofibroblasts enriched for cell–matrix interaction programs. Menopause also enhances homotypic communication within macrophage and myofibroblast subsets, suggesting self-reinforcing loops that may amplify local inflammatory and fibrotic remodeling. **E**, Schematic summarizing the functional reorganization of immune-stromal networks in menopause. Estrogen-responsive Nrip1^+^ macrophages and activated Npnt^+^ myofibroblasts undergo coordinated reorganization in response to menopausal hormonal shifts, forming dynamic heterotypic and homotypic signaling axes that may contribute to fibrotic tissue remodeling. *Abbreviation: HSC/MPP, hematopoietic stem and multipotent progenitor*.

Visualization of cross-tissue interaction networks revealed a subset of preserved edges that represent shared, menopause-associated intercellular interactions across tissues (**Figure 3B**). Our analysis further revealed that transcriptionally distinct subsets of naïve B cells, hematopoietic stem and multipotent progenitor (HSC/MPP), and macrophages consistently emerged as conserved signaling hubs across menopausal tissues (**Figure 3C**). Notably, naïve B cells exhibited transcriptional programs indicative of signaling activity related to fibroblast activation and proliferation, as determined by our iNMF-based analysis (**Figure 3C**), supporting their likely predominance in menopause-associated fibrosis possibly through immune-stromal interaction. Of note, these cells type were assigned using the ScType pipeline, which integrates manually curated positive and negative marker sets to score and annotate cell types.^17^ As an example, the HSC/MPP cluster exhibited high expression of canonical progenitor markers such as *Sca-1* and *Cd34*, supporting their classification as undifferentiated hematopoietic populations.

To further investigate the immune-stromal axis, we focused on macrophages, which expanded in number following menopause across all tissues investigated (**Figure 1C**). Given the well-characterized process of macrophage-to-myofibroblast transition (MMT)—whereby macrophages adopt fibrogenic phenotypes and directly contribute to tissue fibrosis in organs such as the kidney, lung, and liver^23^—we next wanted to know if a similar mechanism may underlie menopause-associated fibrotic remodeling. To this end, we extended our immune cell annotation to include myofibroblasts—contractile, matrix-producing cells that represent the end effectors of fibrosis. This enabled a direct evaluation of macrophage–myofibroblast interactions across all four tissues.

Our analysis revealed dynamic and menopause-specific communication between these two cell types. OVX induced a rewiring of heterotypic communication from estrogen-responsive *Nuclear Receptor-Interacting Protein 1* (*Nrip1)*^+^ macrophages to *Nephronectin* (*Npnt)*^+^ myofibroblasts. These *Npnt*^+^ cells were highly enriched for ligands and receptors associated with cell–matrix interactions and emerged as prominent hubs within the inferred communication network, suggesting a coordinated fibrogenic response. Beyond heterotypic interactions—signaling between different cell types—we also observed menopause-associated changes in homotypic signaling within individual cell type, meaning communication occurring among cells of the same type. Hormone deprivation induced a rewiring of macrophage– macrophage communication networks, particularly between *Nrip1*^+^ and *Cd74*^+^ macrophages, as evidenced by inferred signaling edges in the network graph, suggesting enhanced local immune crosstalk (**Figure 3D**). Similarly, *Npnt*^+^ and *Fibronectin-1* (*Fn1)*^+^, myofibroblasts formed a more extensive myofibroblast–myofibroblast communication network under hormone-deprived conditions, consistent with amplified matrix remodeling (**Figure 3D**). Together, these findings suggest that hormone deprivation drives a functional reorganization of both heterotypic and homotypic signaling axes, enabling hormonally sensitive macrophages and myofibroblasts to adopt new communication patterns during fibrogenesis (**Figure 3E**).

### Tissue-specific cell-cell interactions with menopause

Finally, to dissect tissue-specific intercellular interactions, we applied our iNMF-based cell–cell interaction analysis to each tissue individually with the goal of identifying differential immune responses that contribute to menopausal remodeling (**Figure 4A**). Our analysis revealed that different tissues display distinct immune cell types that serve as central hubs of cell–cell interactions, highlighting tissue-specific immune adaptations to menopause (**Figure 4B**). Specifically, in the liver, endothelial cells and memory CD8^+^ T cells were prominently involved in modulating cell–extracellular matrix interactions. In the lung, HSC/MPP cells acted as key initiators of menopausal tissue remodeling and angiogenesis. In the pancreas, macrophages assumed a central role in orchestrating changes to the local immune landscape. Finally, in skeletal muscle, natural killer cells and macrophages contributed to the regulation of muscle system processes and T cell proliferation (**Figure 4B**). We found our observations in skeletal muscle particularly interesting because our analysis revealed a unique and previously underappreciated role for *Myom2*^+^ natural killer cells in regulating muscle remodeling during menopause (**Figure 4B**). *Myom2* has previously been identified in natural killer cell subset in single-cell transcriptomics in tuberculosis.^24^ While broadly conserved immune responses were observed across multiple tissues, natural killer cell interactions displayed a muscle-specific signature, distinguishing them from other immune cell activities. These findings highlight natural killer cells as potential tissue-specific modulators of menopause-driven skeletal muscle remodeling.

**Figure 4.**
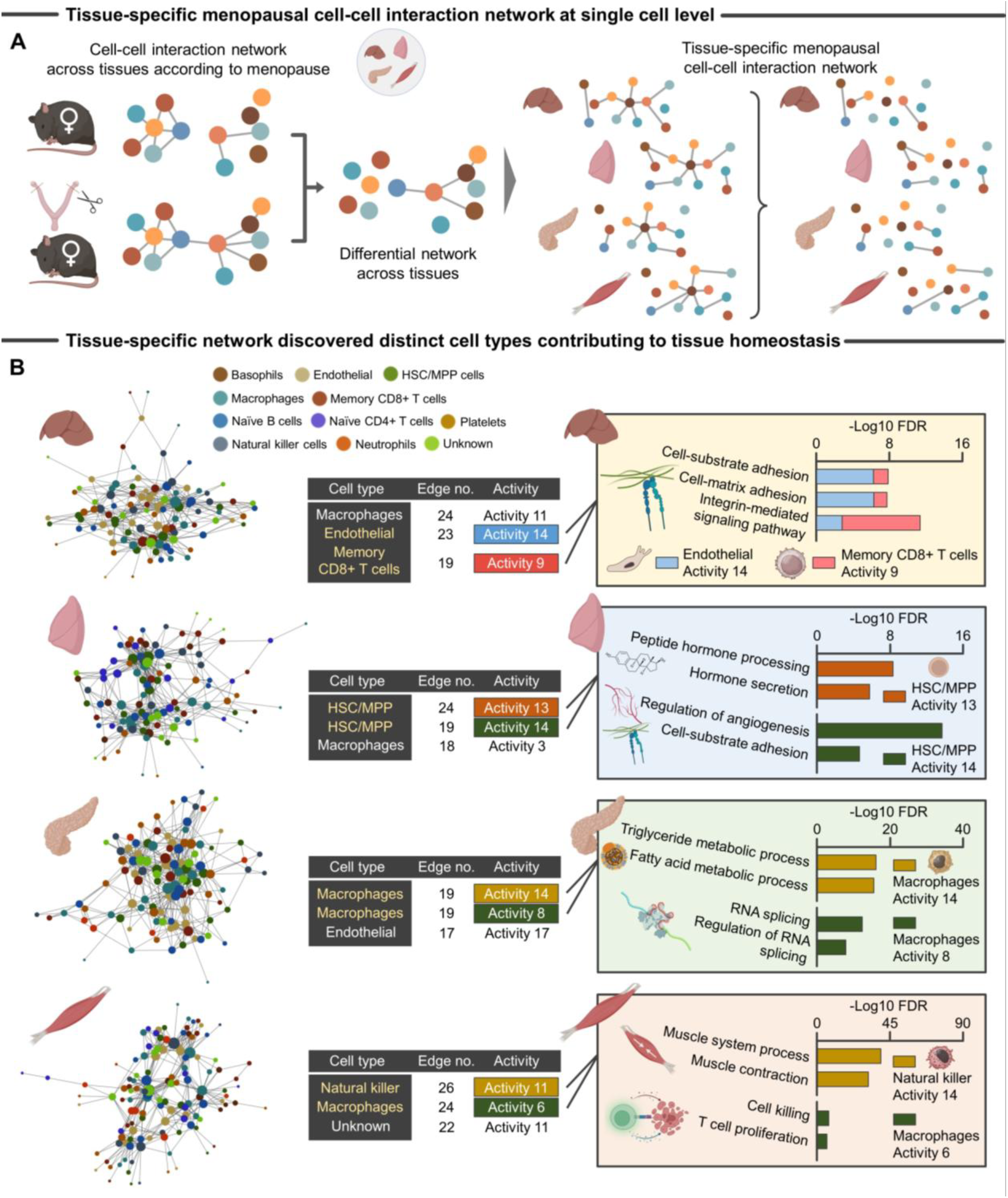
Tissue-specific immune network remodeling in menopause. **A**, Schematic representation of the differential cell-cell interaction analysis framework developed to identify menopause-associated *tissue-specific* shifts in transcriptionally defined cell-cell signaling across four tissues. **B**, Network visualizations of tissue-specific cell-cell interaction networks in liver, lung, pancreas, and skeletal muscle, illustrating distinct menopausal remodeling patterns within each tissue. Key immune populations—such as liver-resident memory CD8^+^ T cells, lung HSC/MPPs, pancreatic macrophages, and skeletal muscle natural killer cells—emerged as tissue-specific signaling hubs. *Abbreviation: HSC/MPP, hematopoietic stem and multipotent progenitor*.

## DISCUSSION

While it has long been recognized in the clinic that menopause induces a systemic remodeling of tissue microenvironments, tissue-specific versus tissue-conserved mechanisms underlying such remodeling remain unknown. Using network inference analysis on scRNA-seq data, we provided a comprehensive, cross-tissue analysis of cell–cell interaction networks in the context of menopausal transition, uncovering a previously unrecognized layer of immune–stromal coordination. By leveraging an integrative network-based framework across four biologically distinct tissues, we discovered that menopause triggers a conserved rewiring of intercellular signaling. Notably, these interaction networks exhibit scale-free topologies, highlighting the presence of highly connected signaling hubs that act as central conduits for propagating menopausal signals systemically. These findings suggest a subset of immune-mediated signaling that serves as a unifying mechanism of menopausal tissue remodeling, providing new insights into how endocrine transitions can shape multicellular dynamics systemically.

We found that the immune system, particularly B cells, HSC/MPP, and macrophages, as central mediators of the coordinated tissue remodeling that occurs during menopause. These cell types consistently emerged as conserved signaling hubs across multiple organs, indicating the existence of shared immune programs that transcend tissue-specific contexts. While previous studies from transgenic mice models have demonstrated an alteration in tissue-resident macrophage population in specific tissues (e.g., brain, skin, and lung) following estrogen receptor deficiency,^25^ our integrative analysis demonstrates that this phenomenon may not be organ-limited but rather may represent a systemic feature of menopausal remodeling. Notably, the identification of naïve B cells as active participants in fibroblast activation and proliferation aligns with recent reports implicating tissue-resident B cells in skin and lung fibrosis.^26^ This study further expands on those findings by showing that naïve B cells directly engage in signaling interactions with fibroblasts across multiple tissues during the menopausal transition. This observation not only supports the emerging view that adaptive immune cells orchestrate stromal reprogramming during endocrine aging but also suggests that conserved immune–stromal crosstalk may represent a unifying mechanism linking hormonal decline to fibrogenic responses across diverse tissues—a hypothesis that may have broad implications for understanding and targeting fibrosis in postmenopausal conditions.

The role of macrophages in this context appears to be particularly multifaceted. Beyond their numerical expansion, transcriptionally distinct Nrip1^+^ macrophages exhibited enhanced interactions with Npnt^+^ myofibroblast subsets, which are characterized by a high expression of genes involved in matrix-related signaling pathways. Nrip1, a nuclear receptor corepressor, is associated with inflammatory activation as a co-activator for NK-kappa B signal in macrophages, while Npnt is a gene that codes for nephronectin, an extracellular matrix protein implicated in fibrotic remodeling.^27,28^ This interaction may reflect a menopause-specific adaptation of the macrophage-to-myofibroblast transition (MMT), a process previously characterized in fibrotic conditions of the kidney, lung, and liver, where macrophages adopt a myofibroblast-like phenotype and contribute directly to extracellular matrix deposition and tissue stiffening.^29-31^ While MMT has traditionally been studied in the context of pathological fibrosis, our findings suggest that a hormonally regulated variant of this process may also occur during physiological endocrine aging. In addition to this heterotypic interaction, we observed strengthened macrophage–macrophage signaling loops, consistent with self-reinforcing inflammatory circuits reported in chronic disease settings.^32^ Notably, these immune-stromal interactions were conserved across multiple tissues, implicating macrophages as systemic mediators of fibrogenic remodeling in response to hormonal decline. These findings position the macrophage–myofibroblast axis as a potential therapeutic target for mitigating multi-organ fibrosis in postmenopausal individuals.

On the other end of the spectrum, we identified tissue-specific adaptations, adaptations we posit reflect the distinct microenvironments and biological demands of each organ. In the lung, HSC/MPP cells emerged as key modulators of menopausal remodeling, possibly via roles in angiogenesis, immune reconstitution, and local hematopoiesis. This aligns with recent reports showing that adult human lungs harbor CD34^+^ hematopoietic progenitors in the alveolar interstitium with immune-enriched gene signatures, suggesting a previously underappreciated extramedullary role during hormonal transitions.^33^ In skeletal muscle, *Myom2*^+^ natural killer cells emerged as tissue-specific hubs. Although *Myom2*^+^ natural killer cells have been identified as driving fibrosis in the setting of tuberculosis where they exhibit diminished cytotoxicity^24^, their role in skeletal muscle and endocrine aging remains unexplored. Functional validation and mechanistic studies will be essential to elucidate how these immune circuits are regulated and whether they can be modulated for therapeutic benefit.

While our study has effectively characterized immune cell rewiring across diverse tissues during the menopausal transition, several limitations should be acknowledged. First, while the OVX model remains a widely used and clinically relevant approach for studying menopausal transitions, it does not fully recapitulate the gradual and complex hormonal changes that occur during natural menopause. That is, the abrupt loss of ovarian hormones in OVX models may exaggerate certain immune or stromal responses compared to endocrine aging observed in humans. Second, our analyses were conducted using relatively young animals at the time of OVX, which may not capture the interplay between hormonal decline and aging-related changes of the tissue microenvironment. Since menopause in humans is typically accompanied by accumulated cellular senescence and age-related tissue remodeling, future studies incorporating aged models will be essential to fully elucidate the synergistic effects of endocrine and chronological aging. Third, although we performed a comprehensive cross-tissue analysis, the scope of our investigation was limited to four organs, excluding other tissues such as bone and tendon that are also profoundly impacted by menopause. Expanding the range of tissues analyzed could further enhance our understanding of systemic remodeling during menopausal transition. Finally, while network-based predictions identified key immune–stromal interactions, functional validation such as lineage-tracing, loss-of-function studies will be essential to establish causal relationships. Despite these limitations, our findings provide a foundational framework for understanding conserved immune–stromal networks underlying menopausal tissue remodeling.

Together, our study provides a comprehensive, cross-tissue map of cell–cell communication in the context of menopause. It reveals how hormonal cues reshape the immune–stromal landscape, highlighting conserved mechanisms that drive fibrotic remodeling while uncovering tissue-specific nuances. These insights not only enhance our understanding of menopause as a systemic biological transition but also provide a foundation for identifying therapeutic targets aimed at mitigating fibrosis, preserving tissue function and promoting healthy aging in postmenopausal individuals.

## MATERIALS and METHODS

### Preprocessing of single-cell RNA-seq data

We accessed archived single-cell RNA sequencing (scRNA-seq) data of skeletal muscle from 14-week-old C57BL/6 female mice that underwent ovariectomy (OVX) or were left untreated (naïve control) at 10 weeks of age. This dataset is publicly available under accession number OMIX001083.^12^ We processed scRNA-seq data using the Seurat R package (version 5.2.0). For each sample, we loaded the gene-barcode matrices from .h5 files and created Seurat objects, filtering cells based on quality control thresholds (genes detected between 200 and 6000; mitochondrial gene content <25%). SCTransform normalization was performed using the glmGamPoi method for variance stabilization and to mitigate sequencing depth differences.

To integrate samples across organs (liver, lung, pancreas, and skeletal muscle), we selected 3,000 integration features and used reciprocal principal component analysis (RPCA) to identify integration anchors. These anchors were then used to perform SCT-based integration, followed by principal component analysis (PCA) and uniform manifold approximation and projection (UMAP) for dimensionality reduction. Clusters were identified using shared nearest neighbor (SNN) graph-based clustering (resolution = 0.5), and cluster-specific marker genes were identified using a likelihood ratio test implemented in Seurat.

For cell type annotation of immune cells, we applied the ScType framework using curated positive and negative marker gene sets from a modified version of the ScType database.^17^ ScType scores were computed based on the integrated expression matrix, and the highest-scoring cell type was assigned to each cluster unless scores fell below one-quarter of total cells in the cluster, in which case the label “Unknown” was assigned.^17^ Representative marker genes were manually reviewed for key immune populations to ensure annotation fidelity—for example, *Cd68, Cd14*, and *Cd11b* for macrophages.

To assess changes in the relative abundance of immune cell populations across experimental groups, we quantified the proportion of each annotated cell type within individual samples. Specifically, we grouped cells by immune cell annotations and calculated the proportion of each immune cell type within each sample. These proportions were then averaged within experimental groups (control and OVX) to identify cell types with higher or lower representation in the OVX group. Macrophages were highlighted in the resulting bar plots to facilitate visualization of relevant shifts in immune composition. All cell type proportion values and groupwise comparisons were exported for further interpretation.

### Transcriptional programs identification in each cell type

Transcriptional programs in each cell type were identified using consensus non-negative matrix factorization (iNMF) across a range of K values, utilizing the LIGER R package (https://github.com/welch-lab/liger) to optimize the number of identified programs.^21^ iNMF is a matrix factorization technique that extends traditional non-negative matrix factorization (NMF) by enabling integration across multiple datasets, allowing the decomposition of gene expression profiles into a set of shared and dataset-specific metagenes. This makes it especially useful for analyzing heterogeneous single-cell RNA-seq data while accounting for batch effects or biological variability across conditions. Metadata from the combined Seurat object was extracted, and the Seurat object was updated to ensure compatibility with the iNMF workflow. The iNMF analysis was then performed across a range of K values, with a maximum K set to 20, using the iNMF_ksweep function. The iNMF_ksweep function systematically explores different K values and outputs consensus factors that are reproducibly identified across multiple iterations, enhancing robustness in factor selection.

The results for each cell type were processed by separating the consensus results (i.e., stable transcriptional programs) and the scaled gene expression data. To assess how transcriptional programs diverge or converge with increasing resolution (higher K values), phylogenetic trees were constructed for each cell type using the build_phylo_tree function. These trees illustrate the hierarchical relationships between identified transcriptional programs and help in evaluating the granularity of the selected K value. This integrative and hierarchical approach enables a more nuanced understanding of cell-type–specific transcriptional landscapes.

To refine the analysis further, potential outlier activity programs—those representing rare contaminating cells—were identified using the identify_outlier_programs function. The distance threshold for partitioning the phylogenetic trees was then determined using the suggest_dist_thresh function. The highest suggested threshold was selected and rounded to the nearest integer for partitioning. Using this threshold, phylogenetic trees were partitioned with the partition_phylo_tree function, based on both the distance threshold and an outlier score threshold. This partitioning allowed for the identification of distinct groups of activity programs for each cell type. These partitions were visualized by plotting the phylogenetic trees with their corresponding activity program groups using the plot_phylo_trees function.

To assess the optimal number of activity programs, the weighted number of subtrees for each choice of NMF rank (K) was calculated using the calculate_K_metric function. The suggested K values were determined by analyzing the weighted subtrees and identifying the K value that maximized this metric. These K metrics were visualized by plotting the weighted number of subtrees as a function of NMF rank K. The optimal K values were marked on these plots, providing a clear indication of the most relevant activity programs for each cell type. Finally, the NMF results for the selected optimal K were obtained using the NMF_results_opt_k function, ensuring the identified activity programs were biologically meaningful and reflected the most relevant clusters for each cell type.

### Cross-tissue menopausal cell-cell interaction network construction

To construct differential cell-cell interaction network, the function to compare networks and extract edges that are different between two networks was first defined. This function takes two network matrices as input and identifies edges present in one network but absent in the other. The differing edges are extracted based on matrix indices and converted into node labels, which are then returned as output. When differences in edges between networks are found, the information about the differing edges is returned as a data frame. Next, networks were compared between OVX and control according to tissue, extracting the differing edges using the previously defined function. Specifically, condition pairs were compared by retrieving networks for each condition, and the function was applied to identify differing edges. This process resulted in the identification of tissue-specific differing edges, which were stored accordingly. For each tissue, common edges across tissues and tissue-specific edges were identified. Common edges were those that appeared across multiple tissues, while tissue-specific edges were those that appeared only in a particular tissue. The edge information was then converted into matrix indices to prepare for network visualization Network visualization was performed using the graph package, where nodes were colored by cell type and edges were displayed in gray.

### Assessment of scale-free network properties

To evaluate the scale-free topology of condition-specific molecular interaction networks, we analyzed the degree distribution (i.e., the number of connections per node) for each node across networks generated under different experimental conditions. Adjacency matrices for each network were constructed based on weighted, undirected interactions without self-loops. These matrices were converted into igraph objects using the graph_from_adjacency_matrix function, enabling the calculation of node degree using the degree function. Each node was annotated by its corresponding cell type and organ. For each condition, we extracted the degree of all nodes and summarized the data by computing the mean, median, and standard deviation of node degrees within each organ–cell type combination. The full summary statistics were exported for downstream interpretation. To visualize the scale-free nature of the networks, we generated density plots of node degrees for each condition, stratified by cell type. Additionally, violin plots were used to compare degree distributions between experimental groups (control versus OVX) within each organ and cell type.

### Network hub analysis

The degree (i.e., the number of edges) for each node in the network was calculated, and nodes were sorted based on their degree. This process identified the nodes with the highest number of edges in the network. The results of the degree-based sorting were saved as a text file and used as part of the network feature analysis.

### Module-specific pathway enrichment analysis

To extract top genes for each module, top genes were retrieved from the results of non-negative matrix factorization (NMF) and listed for each module. The gene symbols were then converted into Entrez IDs. Gene Ontology (GO) biological process (BP) enrichment analysis was performed for the top genes in each module, identifying significant pathways. The results of the enrichment analysis were stored as lists for each cell type and module, and these results were then merged into a single data frame. The final enrichment results were saved as a text file and used for downstream analysis.

### Pseudo-bulk differential gene expression analysis

To identify genes that are differentially expressed between control and OVX groups within macrophages, we constructed pseudo-bulk expression profiles by summing the raw read counts of all single macrophage cells from each sample. Only samples with at least 100 macrophages were retained to ensure sufficient cell representation. To mitigate differences in sequencing depth, each pseudo-bulk sample was randomly down-sampled to match the minimum library size observed across all included samples. Genes were filtered to retain only those with a raw count of at least 10 in all samples, ensuring consistent expression across the dataset. Differential gene expression analysis was performed using the DESeq2 R package with group as the design factor.^34^ Log_2_ fold changes were computed as OVX versus control (i.e., log_2_ (OVX/control)). P-values were adjusted using the Benjamini-Hochberg method to control the false discovery rate.

To explore the overlap of differentially expressed genes across experimental groups, Venn diagrams were constructed using genes that satisfied the threshold of |log_2_(fold change)| ≥ 1 and adjusted *p*-value < 0.05. For each comparison, gene lists were extracted from the DESeq2 results, and file names were parsed to label the groups accordingly. Diagrams were generated using the VennDiagram R package.

### Statistical analysis

All statistical analyses were performed using R studio software RStudio (Posit PBC, Boston, MA). Except where indicated, data are displayed as means, with uncertainty expressed as 95% confidence intervals. For unpaired experiments, two-tailed Student *t*-test was performed. We checked the features of the regression model by comparing the residuals vs. fitted values (i.e., the residuals had to be normally distributed around zero) and independence between observations. No correction was applied for multiple comparison because outcomes were determined *priori* and were highly correlated. In all experiments, p-values <0.05 were considered statistically significant. Except where indicated, throughout this text, “*n*” represents the number of independent observations. Specific data representation details and statistical procedures are also indicated in the figure legends.

## Acknowledgments

This study was supported in part by (1) NIA R01AG089455 for FA and HI, and (2) NIA R01AG061005 for FA. The funders had no role in study design, data collection and analysis, decision to publish, or preparation of the manuscript.

## Competing interests

None declared.

## Notes

### Competing Interest Statement

The authors have declared no competing interest.

